# Making Common Fund data more findable: Catalyzing a Data Ecosystem

**DOI:** 10.1101/2021.11.05.467504

**Authors:** Amanda L Charbonneau, Arthur Brady, Karl Czajkowski, Jain Aluvathingal, Saranya Canchi, Robert Carter, Kyle Chard, Daniel J.B. Clarke, Heather H Creasy, Mike D’Arcy, Michelle Giglio, Alicia Gingrich, Rayna M Harris, Theresa K Hodges, Olukemi Ifeonu, Minji Jeon, Eryk Kropiwnicki, Marisa C.W. Lim, R. Lee Liming, Meisha Mandal, James B Munro, Suvarna Nadendla, Rudyard Richter, Cia Romano, Philippe Rocca-Serra, Robert E. Schuler, Hongsuda Tangmunarunkit, Alex Waldrop, Cris Williams, Karen Word, Susanna-Assunta Sansone, Avi Ma’ayan, Rick Wagner, Ian Foster, Carl Kesselman, C. Titus Brown, Owen White

## Abstract

The Common Fund Data Ecosystem (CFDE) has created a flexible system of data federation that enables users to discover datasets from across the U.S. National Institutes of Health Common Fund without requiring that data owners move, reformat, or rehost those data. The CFDE’s federation system is centered on a catalog that ingests metadata from individual Common Fund Program’s Data Coordination Centers (DCCs) into a uniform metadata model that can then be indexed and searched from a centralized portal. This uniform Crosscut Metadata Model (C2M2) supports the wide variety of data types and metadata terms used by the individual DCCs and is designed to enable easy expansion to accommodate new data types. We describe its use to ingest and index data from ten DCCs.

## Introduction

Findability of existing data in biomedical research is important for data reuse. Reusing data can increase the speed of scientific discovery, as well as allow researchers to generate hypotheses [1–3]. However, these benefits are highly dependent on the Findability, Accessibility, Interoperability, and Reusability (FAIRness) [4] of the individual datasets, with Findability by interested researchers being the first essential step. Improving reuse of datasets is increasingly a priority for both scientists and funding agencies [5–7], and there are many data publishing and scientific data repositories that are designed to enable targeted search across a breadth of data types [8]. However, many researchers find it difficult to navigate the many available data repositories or to predict which might hold data that are useful to them [9]. Further, researchers browsing a data repository typically rely on topical information to determine if it contains relevant data [10–12], and as most large data repositories typically do not impose constraints on terms used to describe datasets, they are not always well suited to browsing [8].

The U.S. National Institutes of Health (NIH) Common Fund (CF) was created in 2006 to fund biomedical research efforts that did not fit into the funding remit of any one NIH Institute or Center, with the intention of generating unique and catalytic datasets that would serve as resources to fuel future innovation. Nearly fifteen years later, more than fifty CF Programs have been funded and have created large, diverse collections of genomic, transcriptomic, proteomic, metagenomic, and imaging assets. These data are deep, derived from hundreds of studies, with samples collected from thousands of human subjects, cell lines, organoids, and animal models. Each CF Program has a Data Coordination Center (DCC) which facilitates program data storage in repositories or local data centers for public use, and most also host curated, derived datasets at program-specific data portals to enable easy use by biomedical researchers. For example, the Genotype-Tissue Expression (GTEx) project [13] offers sophisticated search and contextual display of processed gene expression, tissue characteristics, and quantitative trait loci (QTLs) at their web portal. The GTEx data portal sees over 15,000 users a month [14], and has enabled hundreds of published studies where the GTEx data was reused by the broad research community.

Although the Common Fund was created to catalyze cross-cutting research and create reusable datasets for biomedical research [15], the many datasets created by different CF Programs are not located in a single repository or accessible via a common interface. This is an artifact of the Common Fund funding model. As of 2019 individual CF projects were isolated, with few connections between active projects, and there were few incentives to integrate them [16]; rather, each of the Programs had created its own metadata, storage, and access solution. This proliferation of access and storage methods is especially problematic for biomedical researchers, who show a strong interest in reusing data, and are the target user base for Common Fund, but generally lack expertise in using data repositories [6]. The Common Fund Data Ecosystem (CFDE) was established in 2019 to address both the siloed nature of CF data, and the downstream impact this has on biomedical data reuse.

Prior to the CFDE, Common Fund-funded programs typically had little or no contact, let alone active collaboration, and with no links between portals it was challenging for a researcher to navigate across Common Fund resources. This independence and isolation of different Common Fund programs has benefits, in that it allows each program to tailor their data gathering, portals, and infrastructure to answer domain-specific questions, and to respond nimbly to changes in program needs. However, this independence has also impeded data integration around common data types. Even the seemingly simple task of finding what data are available is hindered by differences in nomenclature.

Comparing data across programs is particularly challenging. Each program’s data portal provides a curated experience of analyzed data that usually does not support comparison with data from other sources. Moreover, many data can only be meaningfully compared to other data analyzed in the same way, and as each Common Fund program operates independently, data are stored, labeled, analyzed, curated and maintained in incompatible ways. Thus, a researcher interested in combining data across CF programs is faced with not only a huge volume, richness, and complexity of data, but also a wide variety, richness, and complexity of data access systems with their own vocabularies, file types, and data structures. Reusing Common Fund data for new cross-cutting analysis requires expertise in working with large files, accessing data in the cloud, harmonization, and data transformation -- all before any scientific analysis can begin. Each stage presents an *individually* large challenge for a typical biomedical researcher or clinician which motivates labs to hire dedicated bioinformaticians (at considerable cost to NIH); confronting all of these challenges together is prohibitive for integrative analyses.

To make Common Fund data more findable, the CFDE has created a flexible system of data federation that enables users to discover datasets from across the CF at a centralized portal without requiring Common Fund programs to move, reformat, or rehost their data, similar to the federation strategy of the Research Data Alliance [17], The Australian Research Data Commons [18], and the Earth System Grid Federation [19]. The CFDE uses a sociotechnical federation system that combines proven, explicitly community driven approaches [17,20,21] with a model-driven catalog that incorporates metadata submitted by individual CF Program Data Coordination Centers (DCCs) into a uniform metadata model that can then be indexed and searched from a centralized portal.

The sociotechnical framework of the CFDE is a self-sustaining community that both harmonizes existing data and develops community standards that newly funded programs can use to interoperate with existing datasets. Governance of the CFDE includes extensive use of “cross-pollination” networking events, Requests for Comments (RFC) documents for community input, and documentation of use cases. In addition, working groups have been established to guide best practices in areas such as gene-centric knowledge representation, clinical metadata, ontologies, technical implementation strategies, and genetic variants. All of this community effort manifests itself as the CFDE Portal, a single user-friendly search interface for the Common Fund where all data is searchable using a common model. This uniform *Crosscut Metadata Model* (C2M2) supports the wide variety of dataset types, vocabularies, and metadata terms used by the individual CF DCCs. This C2M2 is designed to enable easy expansion to accommodate new data types that may emerge from existing DCCs or as new DCCs join the CFDE, and as new use cases are embraced.

The primary user interface for the CFDE’s metadata catalog is a Web-based portal [22] that supports multi-faceted search of metadata concepts such as anatomical location, species, and assay type, across a wide variety of datasets using controlled vocabularies. This style of search supports common researcher use cases [10,11] by giving them the ability to filter data based on their desired topical information to more easily discover datasets that would otherwise require idiosyncratic targeted search across multiple databases. Findability is more than just search, it is the user’s experience of interacting with the search and understanding how to use it to find relevant results. We work closely with a professional usability testing team to ensure that our portal meets user needs.

In this paper, we describe the motivation for the C2M2, detail the current C2M2, and discuss the portal that serves gathered metadata from across CF programs. We also describe the strategy that guides C2M2 development and the processes by which the C2M2 evolves.

## Results

### Common Fund data cannot be found by uniform internet search terms

A hypothetical biomedical researcher interested in finding Common Fund RNAseq datasets created from human blood samples should, in theory, be able to find relevant data from at least five Common Fund programs: Genotype-Tissue Expression (GTEx), Gabriella Miller Kids First (GMKF), Human Microbiome Project (HMP), Extracellular RNA (ExRNA), and the Library of Integrated Network-Based Cellular Signatures (LINCS). Each of these programs hosts their data on a public website, and typically also have informational websites about their work, and so would be expected to be easily Findable. However, a Google search for ‘human blood RNAseq “common fund”’ returns 20,500 results, all but 55 of which are omitted by Google as they are “very similar to the 55 already displayed” [23]. These 55 results contain references to data from only three Common Fund programs: GTEx, Human BioMolecular Atlas Project (HuBMAP), and Illuminating the Druggable Genome (IDG). Of these three results, GTEx is in fact the only Common Fund program with RNAseq data from human blood samples. HuBMAP does not have data from blood samples, but Dr. Phil Blood is the director of HuBMAP, so his name matches the search. IDG also lacks RNAseq data from blood, but does feature a blog post that mentions both RNAseq and blood separately. The other four expected programs, GMKF, HMP, ExRNA and LINCS, do not appear in the results.

To illustrate why so few relevant data appear in these search results, we chose six concepts that are broadly applicable to biomedical data -- Sample Type, General Tissue, Specific Tissue, Anatomical Part, Analysis Pipeline, and Organism -- and used them to manually search the five Common Fund RNAseq datasets that we know have RNAseq data from human blood. We then documented how each Common Fund program described these concepts in their respective data portals.

The example search in Figure 1 highlights several common types of differences between Programs. For each of the six concepts, each sub-table lists the “key” used by each Program to refer to that idea, which is equivalent to the column name in a spreadsheet. The “value” is an example of the data you might find under that column, and here we display the values that best fit our “human blood RNAseq” search. These results show three general types of disagreement in term use: differing term values, differing keys in specific categories, and differing assumptions. Differing term values (“RNAseq” vs “RNA-seq” vs “RNA-Seq”) hinder search because the term of interest may not be matched by a search engine.

**Figure 1:**
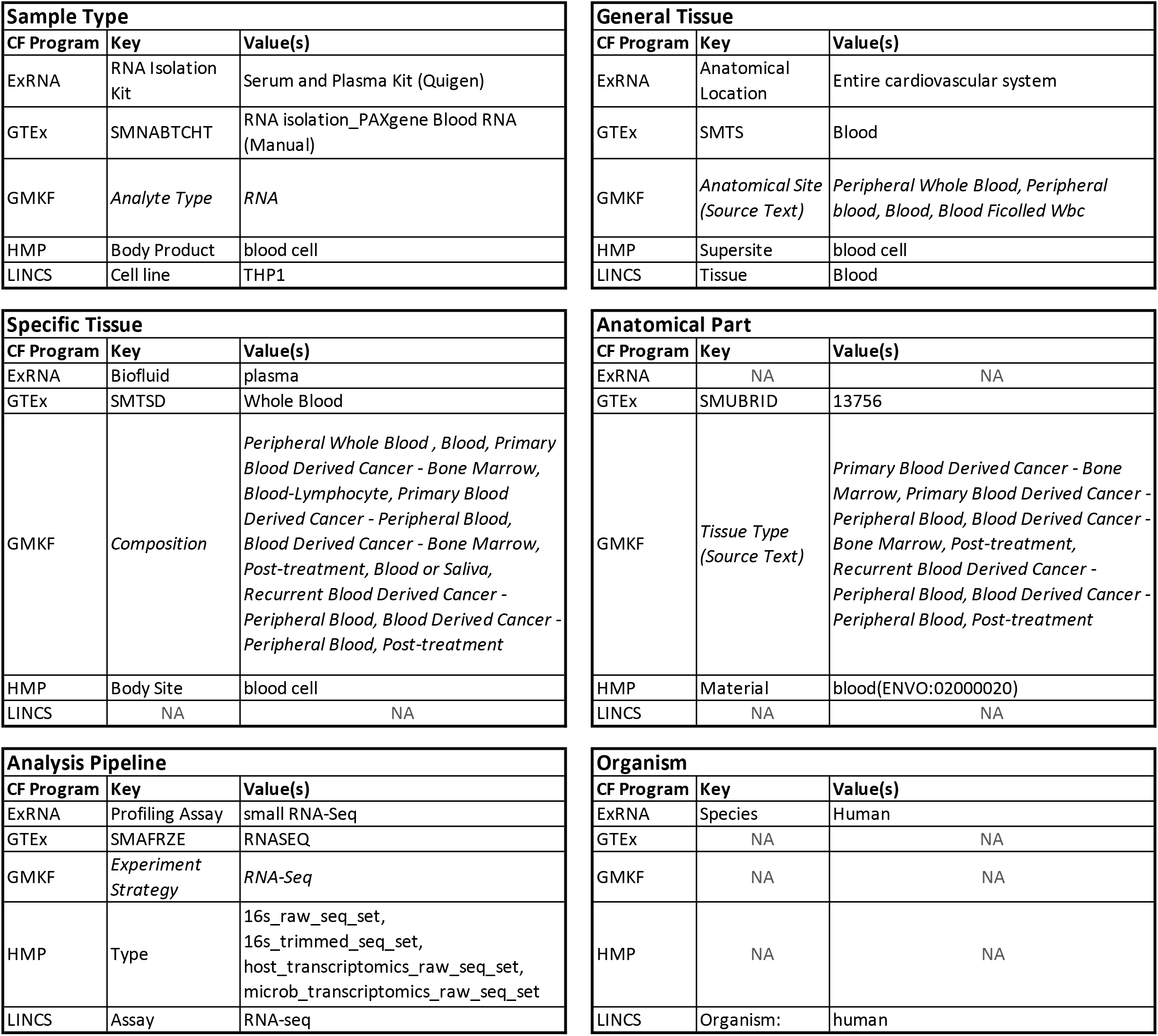
For each of the six concepts (Sample Type, General Tissue, Specific Tissue, Anatomical Part, Analysis Pipeline, Organism) that would be relevant for finding existing datasets for “human blood RNAseq”, we list the Key and Value used by each of five Common Fund programs that host this type of data in their search portals. Keys are analogous to column headers in a metadata file, and the values shown are the specific values used at that program that are good matches for this search. NAs indicate that information for that concept is not an available search term at a given portal. GMKF Keys and Values shown as italics denote that while those terms are publicly available, they can only be searched while logged into the GMKF portal, and so do not appear in Google searches.

Differing keys rarely impact search, but introduce complexity in combining datasets, as each key must be manually harmonized. Figure 1 shows one method of harmonization, but there are other valid interpretations. As each Program uses different terms to describe these concepts, and those terms are often not intuitive, a researcher needs deep familiarity with each dataset to make decisions. For example, is HMP’s “Supersite” most analogous to GTEx’s “SMTS”? Or to GTEx’s “SMTSD”? A researcher who wants to combine these data would need to research what all these terms are and how each site uses them. Differing assumptions can be seen in the reporting for Organism. HMP, GTEx and GMKF only host human data, and thus do not specify species in their internal metadata, making them more difficult to find. Taken together, these differences make data discovery nearly impossible with a uniform set of search terms.

### A listening tour identified obstacles to interoperation

We conducted in-depth interviews in 2019 with nine different Programs to better understand the obstacles that the Common Fund Data Coordination Centers (DCCs) face in making CF datasets more accessible to researchers and to learn what data they collect, how they model and store those data, and their target user base [14,16,24]. We used these visits to identify CF program requirements and establish an initial working relationship with DCCs. A primary outcome of the listening tour was a new draft of a common set of data elements that could describe data held at all current DCCs: the C2M2. Over the ensuing two years, we elaborated the C2M2 through a consensus-driven process, instantiated the C2M2 in a rich relational database, used the C2M2 to ingest data from ten DCCs, and built a web-based portal on top of the ingested data.

### Entities and associations are key structural features of the C2M2

We use the term “entities” to denote C2M2 metadata objects that represent resources of experimental interest. C2M2 offers three **core entities** to represent common (although certainly not universal) components of biomedical experiments: **biosample**, (digital) **file**, and **subject** organism. Two more C2M2 entities represent **containers** (sets or groups *containing* particular files, samples and/or subjects): **project**, representing research studies governing the experiments being documented, and **collection**, a generalization of “dataset” that can include representations of samples and subjects in addition to the digital resources (files) typically exclusively comprising a dataset. Entities are represented in C2M2 as **tables** (rectangular matrices), each **row** of which is a **metadata record** comprising a small list of named columns (**fields**) containing metadata values. Each row describes one instance of whatever entity that row’s table represents: a single file or biosample, for example. Each *field* in each row has an agreed-upon meaning that helps to describe the thing being represented by the row as a whole. **Relationships between entities** are represented as **association tables**, whereby metadata records (rows) describing different types of entities are linked to one another according to broad relationship definitions like “file describes biosample” or “biosample from subject.” See Figure 2 for a simplified **entity-relationship (ER) diagram** describing the model; see Figure supplement 1 for the full C2M2 ER diagram.

**Figure 2:**
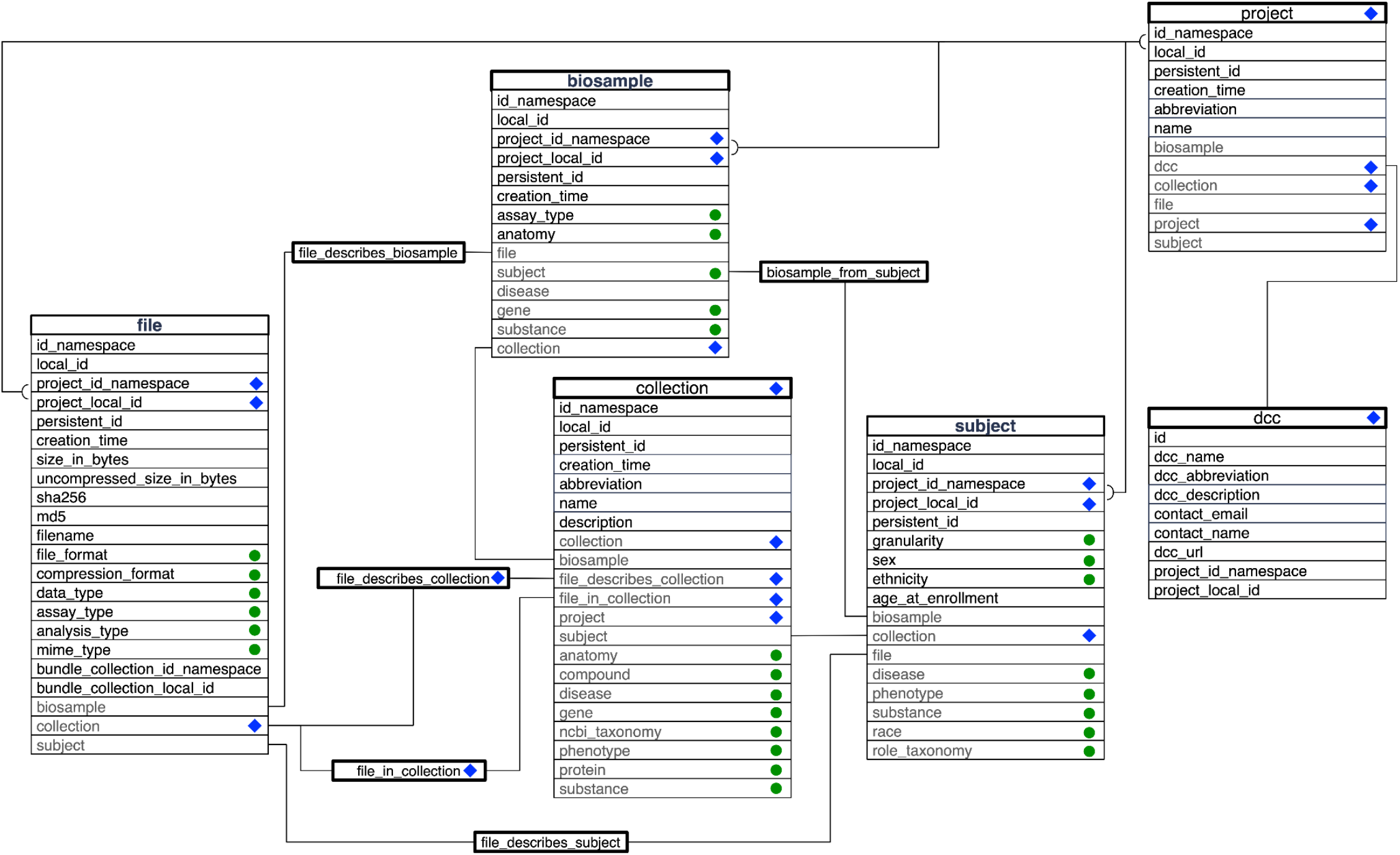
A simplified entity relationship diagram for the C2M2 where most association and controlled vocabulary tables are collapsed into the core and container entity tables. Each core and container entity (file, subject, collection, etc) is shown as a large table. Fields shown in black are directly specified in the table. Fields labeled with blue diamonds indicate that the field functions as a container. Fields marked with green circles indicate that the field must be populated from a controlled vocabulary which is defined in another (not shown) table. Grey fields indicate that the association is made in an association table (not all shown). A grey field with a green circle indicates that the association table also has its own (not shown) controlled vocabulary table. Lines are drawn connecting the fields that participate in the relationship between entities. Named boxes that fall on these paths give the name of the association table that makes the connection, but do not list the tables fields.

As noted above, C2M2 currently supports three core entities describing fundamental types of experimental resources: **files** (digital bytestreams encoding experimental data); **biosamples** (living material collected and processed via experimental protocols); and **subjects** (organisms studied by experiment, both directly observed and as biosample donors). Also available are two “container” entities -- **project** and **collection** - designed to allow DCCs to explicitly and flexibly group related entity records into named sets. A valid C2M2 submission must provide minimal information wherein every C2M2 core entity record (file, biosample, or subject) is linked to exactly one C2M2 project record describing the research effort under which the resource was created or observed. Operations essential to discovery (sorting, searching, and binning) depend on this information, so that as C2M2 resource information is discovered by users, it can be more easily associated with its proper research context. The C2M2 collection entity is a conceptual generalization of “dataset” (a named, well-defined collection of data resources) to also explicitly include non-data resources (like biosamples and subjects). Aggregation of C2M2 records into collections is optional, with decisions defining scope and complexity of usage generally left to the submitting DCC. Collections can optionally be assigned persistent IDs (like DOIs) for stable citation, reference, and retrieval.

A C2M2 record need not (and generally will not) have values for every possible metadata field in order to be usable, especially as the C2M2 broadens to accept new data types and variants: most C2M2 field values are consequently optional. Nearly every column, and most tables, can optionally be left blank, allowing each DCC to build their C2M2 submission with whatever level of richness or focus best fits their capacity and presentation goals, while meeting universal criteria designed to permit basic interoperation and discovery that are kept conservatively minimal by design.

### The C2M2 integrates standardized vocabularies

A key component of cross-DCC metadata harmonization within the CFDE is support for the detailed description of C2M2 metadata using terms from standard scientific ontologies. C2M2 currently provides a variety of features by which controlled (standardized, curated) scientific vocabulary terms can be attached to C2M2 collections and core entities. All C2M2 controlled vocabulary annotations are optional. Currently supported controlled vocabularies, listed in Table 1, include the Disease Ontology [25]; the Ontology for Biomedical Investigations [26]; the Uber-anatomy Ontology (UBERON; [27]; the NCBI Taxonomy [28]; EDAM [29], an ontology for bioinformatics concepts including data types and formats; gene terms from Ensembl [30], a database for researchers studying genomics in humans and other vertebrates and model organisms; PubChem [31] the world’s largest curated cheminformatics database; GlyTouCan[32], a glycan structure repository; the Human Phenotype Ontology[33], a collection of terms describing various external conformations of human anatomy; and UniProtKB[34], the world’s largest collection of protein metadata, combining both expert-curated and auto-annotated information.

**Table 1.** Controlled vocabularies currently supported in C2M2. Entity term fields are listed as `C2M2_entity_table.fieid_name`; term association tables (one-to-many relationships between entities and vocabulary terms) are listed by table name. We give the source ontology for each vocabulary, along with a general description of its annotation role within C2M2.

| CV field or association | ontology | description |
| --- | --- | --- |
| `file.analysis_type` | OBI | the type of analytic procedure that produced a `file` |
| `file.assay_type` | OBI | the type of experiment that produced a `file` |
| `file.file_format` | EDAM | the digital format or encoding of a `file` (e.g. "FASTQ") |
| `file.compression_format` | EDAM | the compression format of a `file` (e.g. "bzip2", "gzip") |
| `file.data_type` | EDAM | the type of information contained in a `file` (e.g. "sequence data") |
| `biosample.assay_type` | OBI | the type of experiment that produced a `biosample` |
| `subject_phenotype` | Human Phenotype Ontology | link `subjects` to phenotypic observations |
| `biosample.anatomy,`<br>`collection_anatomy` | UBERON | the physiological source location in or on a `subject` from which a `biosample` was derived, or an anatomical part relevant to a particular `collection` |
| `biosample_disease,`<br>`subject_disease,`<br>`collection_disease` | Disease Ontology | link `biosamples`, `subjects` and `collections` to observations about diseases |
| `biosample_gene,`<br>`collection_gene` | Ensembl | link `biosamples` and `collections` to individually relevant genes (e.g. knockdown targets) |
| `biosample_substance,`<br>`subject_substance,` | PubChem | link `biosamples`, `subjects` and |
| collection_compound,<br>collection_substance |  | collections to drugs,<br>reagents, other small<br>molecules |
| ncbi_taxonomy.id,<br>subject_role_taxonomy,<br>collection_taxonomy | NCBI Taxonomy | link subjects or<br>collections to taxonomic<br>names |
| collection_protein | UniProt KnowledgeBase | link collections to<br>individually relevant proteins |

If sufficiently specific terms cannot be found in the supported ontologies, we encourage DCC data managers to include, provisionally, more general parent terms (as available), and simultaneously to contact the CFDE Ontology Working Group (WG) with descriptions of any needed additions to the supported controlled vocabularies. CFDE has established direct update channels with the curation authorities for each supported ontology, and the Ontology WG aims to expedite the addition of any missing terms on behalf of Common Fund DCCs. Our search portal directly supports both the official ontologies and the new provisional terms in its pipeline so that DCCs can use the best terms right away rather than resorting to inappropriate term usage.

### The C2M2 supports a flexible system of internal and global identifiers

The C2M2 is designed to be a framework for sharing information with the global research community about data deriving from experimental resources. Experimental metadata are created at different times by different DCCs working independently, so any system trying to federate such information must establish a standard way for DCCs to generate stable identifiers (IDs) without requiring DCCs to coordinate ID usage directly with each other. At the same time, any integrated system must guarantee unambiguous IDs. As such, a minimally effective ID scheme must allow DCCs to create IDs for their C2M2 metadata that do not (and will never) clash with C2M2 IDs created by other (unknown and possibly inaccessible) DCCs.

The C2M2 provides two types of IDs: a mandatory C2M2 ID and an optional persistent ID. C2M2 records for files, biosamples, project, subjects and collections must each be labeled with a C2M2 ID. A C2M2 ID has two parts: a prefix (id_namespace) representing the DCC (i.e., a name identifying the point of origin of the metadata), and a suffix (local_id) representing the specific entity (file, project, etc.) being identified. The two parts of each C2M2 ID, concatenated, serve as a unique ID for each resource that is unambiguous across the entire CDFE ecosystem. This scheme allows DCCs to import their preferred intramural ID scheme directly into the local_id component, generally without modification. The optional persistent ID - stored separately from C2M2 IDs - is a URI that encodes actionable information that users or automated software can follow to get more information about the resource named by the ID, including access or download details, where applicable. The CFDE system accepts a wide variety of domains and URI schemes for persistent IDs, including minids[35], Data Repository Service (DRS) IDs, and digital object identifiers (DOIs); see the Identifiers Supplement for details.

### Independent “data packages” are submitted to the CFDE

The C2M2 is designed to integrate asynchronous submissions from multiple Programs, operating independently. Each submission comes as a “data package”, a collection of data tables encoded as tab-separated value (TSV) files. Each DCC collects metadata for data resources within their Program into a single data package that it then submits to the CFDE. DCCs can explore submitted data packages in advance of publication using protected areas of the CFDE portal; multiple submissions are possible between public data releases, and only the most recent (approved) version of each DCC’s data package will be published as part of each public release.

A C2M2 data package consists of 48 TSV files (as of 4/4/2022) populated with interrelated metadata about DCC data assets. Precise formatting requirements for data packages are specified by a JSON Schema document. This schema is an instance of the Data Package meta-specification published by the Frictionless Data group [36], a platform-agnostic toolkit for defining format and content requirements for files on which automatic validation can then be performed. Using this toolkit, the C2M2 JSON Schema specification defines foreign-key relationships between metadata fields (TSV columns), rules governing missing data, required content types and formats for particular fields, and other constraints. These architectural rules provide guarantees for the internal structural integrity of each C2M2 submission, while also serving as a baseline standard to create compatibility across multiple submissions received from different DCCs.

A data package can be created at several levels of complexity: many columns and several entire tables can be left empty and still produce a valid package for submission. Only three metadata records (three rows, across three C2M2 tables) are strictly required, so most tables can optionally be left empty in a minimally compliant submission. The three required records are:

1. a short contact record (name, email address, and other contact details) referencing the DCC technical contact responsible for the submission;
2. a single project record representing the submitting DCC (for resource attribution); and
3. at least one identifier (ID) namespace, registered in advance with the CFDE, that protects IDs from conflicts with IDs generated by other DCCs.

The simplest *usable* submission configuration will also contain at least one non-empty data table representing a flat inventory of experimental resources (e.g., files, subjects, or biosamples). A more complex submission might inventory additional resource (entity) types, and might also encode basic associative relationships among those entities. For example, a submission might record which biosamples were materially descended from which subjects, or which files contain data pertaining to which biosamples. Beyond the single mandatory project record for resource attribution, DCCs can also construct a hierarchy of project records to subdivide their experimental metadata, grouping resources by funding source (or any other well-defined, “study”-like subdivision of research effort).

### Preparing a project submission for ingest into the catalog

Each TSV file in a C2M2 submission is a plain-text file representing a tabular data matrix, with rows delimited by newlines and fields (columns) delimited by tab characters. Field values in TSV files must conform to all formatting and relational constraints specified in the C2M2 schema document [37]. Any blank table will be represented by a TSV file containing just one tab-separated header line which lists the (empty) table’s field names. Requiring that files exist even for empty tables differentiates intentional data omission from accidental file omission.

For each controlled vocabulary supported by C2M2, a term table must be included as part of any valid submission (see Figure 2, Supplemental Figure 1, green tables). Each such table will contain one row for each (unique) controlled vocabulary term used anywhere in the containing C2M2 submission, along with basic descriptive information for each term that empowers both downstream user searches and automated display interfaces. All term metadata are loaded directly from the ontology reference data: once the C2M2 entity and association tables are prepared, a CFDE-provided script [38] is used to automatically scan the prepared tables for controlled vocabulary terms. These controlled vocabulary terms are validated against externally-provided ontology reference files, and then combined with descriptive data drawn directly from the reference files. The resulting information is then used to automatically build all necessary term tables. These automatically-generated term tables (organized as TSV files) are then bundled along with the rest of the C2M2 submission.

### The CFDE provides a robust data ingest and validation process for data packages

The CFDE Coordination Center provides a full service submission system for data packages. DCCs submit candidate data packages to the CFDE Data Submission System by using the cfde-submit tool, a lightweight command-line Python package [39] that enables authenticated upload to the CFDE Portal via Globus Flows [40] (Figure 3). This tool takes a directory of TSV files (the tables in Figure 2) as input, performs initial validation against the C2M2 model, and builds the directory into a bdbag [41] data package that it securely uploads to a Globus [42] endpoint. Each DCC has a separate, secure Globus endpoint location that is created by the CFDE Coordination Center as part of DCC onboarding, and only authorized DCC users can submit to that DCC’s location.

**FIGURE 3.**
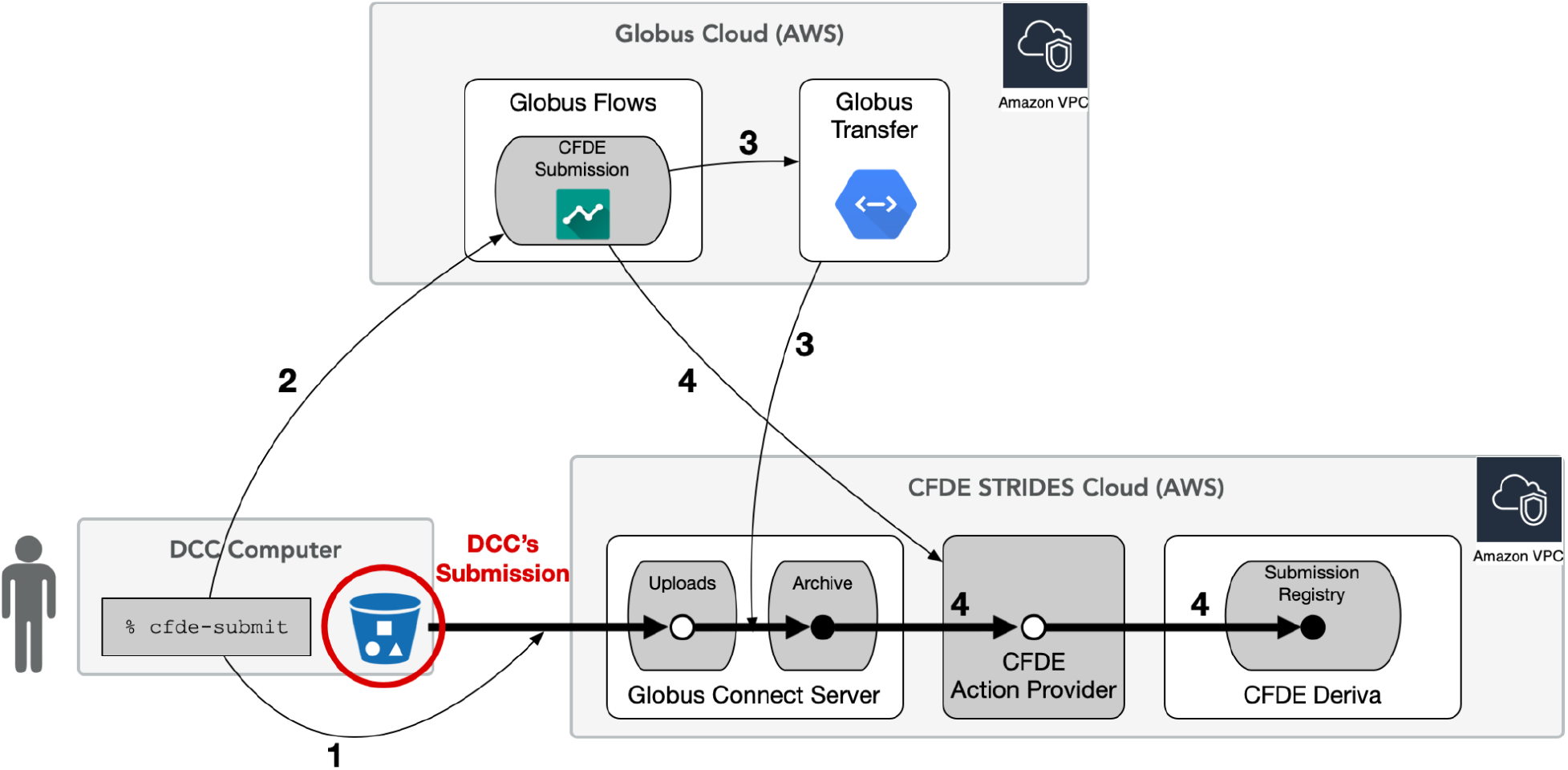
CFDE submission process, focusing on Globus Flows. The `cfde-submit` CLI performs a lightweight validation of the submission data, starts the data upload to CFDE’s servers (step 1), and then initiates processing in the cloud (step 2). The system that manages the cloud processing is called Globus Flows. Globus Flows is Globus software-as-a-service (SaaS) running in the AWS cloud. CFDE’s submission process is one of many “flows” that the Flows service manages, and the final action of `cfde-submit` is to start a run of the CFDE submission flow. The CFDE submission flow moves the submitted data to a permanent location (step 3), sets access permissions (not shown), and executes code on a CFDE server (step 4) that ingests the submitted data into the CFDE Portal’s database service, Deriva. While processing is happening in the cloud (steps 2-3), status can be checked using `cfde-submit`, but it doesn’t appear in the CFDE Portal until step 4.

Once a data package is uploaded to the Globus endpoint, the Deriva [43] database automatically begins ingesting it, performing further validation by using a custom validation script. Depending on the size and complexity of the data package, this process can take up to an hour to complete. Users are notified by email when the submission process has completed, and are provided with a link to view the data or a description of any errors encountered. Processed data packages are viewable by the submitting DCC in secure pages of the CFDE Portal. These pages provide an instance of the exact search pages that will be available to the public if and when the DCC approves the submission, plus a high-level overview of the submission and various summary statistics (Figure 4). Only after the DCC approves the data package is it merged into the CFDE public catalog and made viewable and searchable in the CFDE portal. A high level overview of the entire submission workflow is shown in Figure 5.

**Figure 4:**
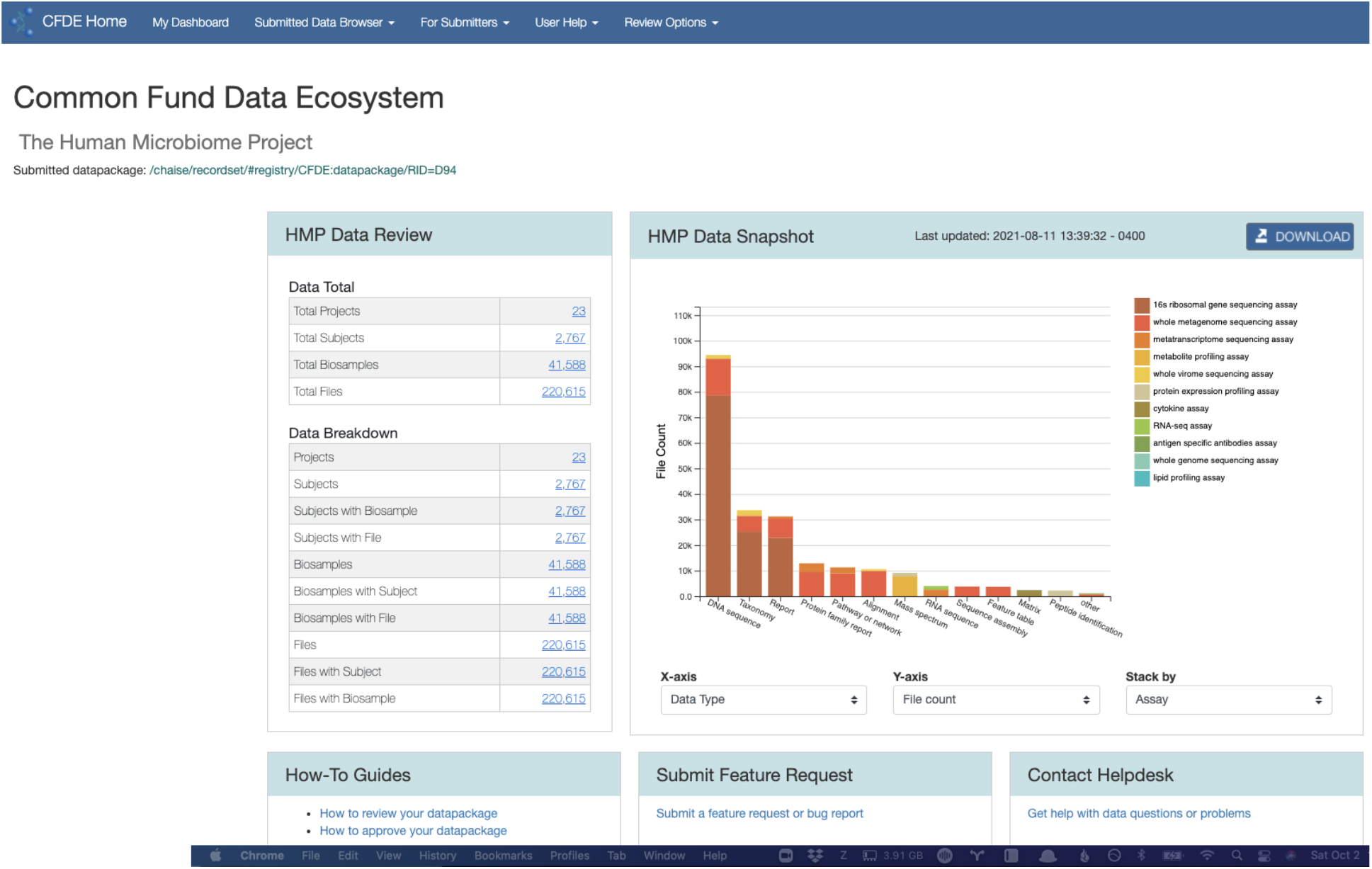
Summary page of a submitted data package with interactive chart, and summary statistics

**Figure 5:**
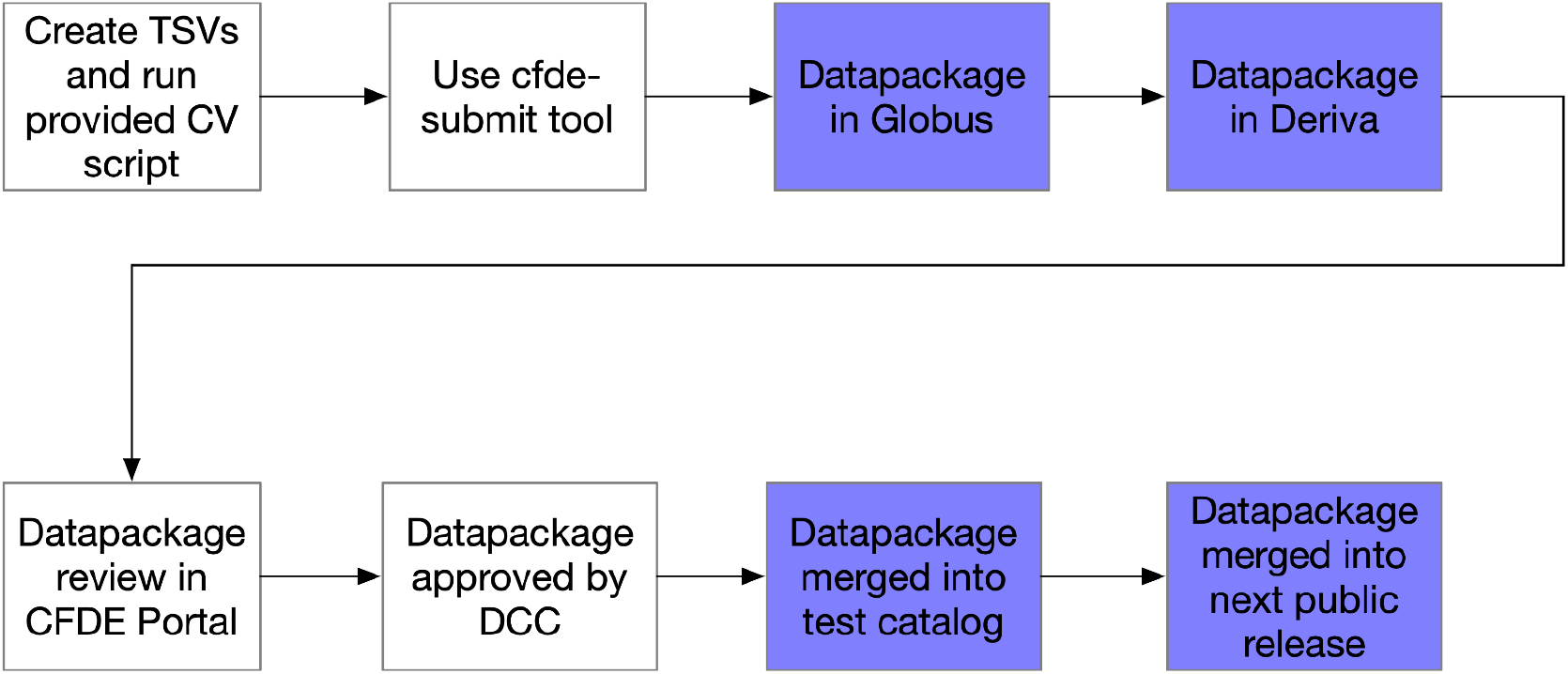
Graphic overview of the steps for data submission. White boxes are user steps, blue boxes are automated.

**Figure 6:**
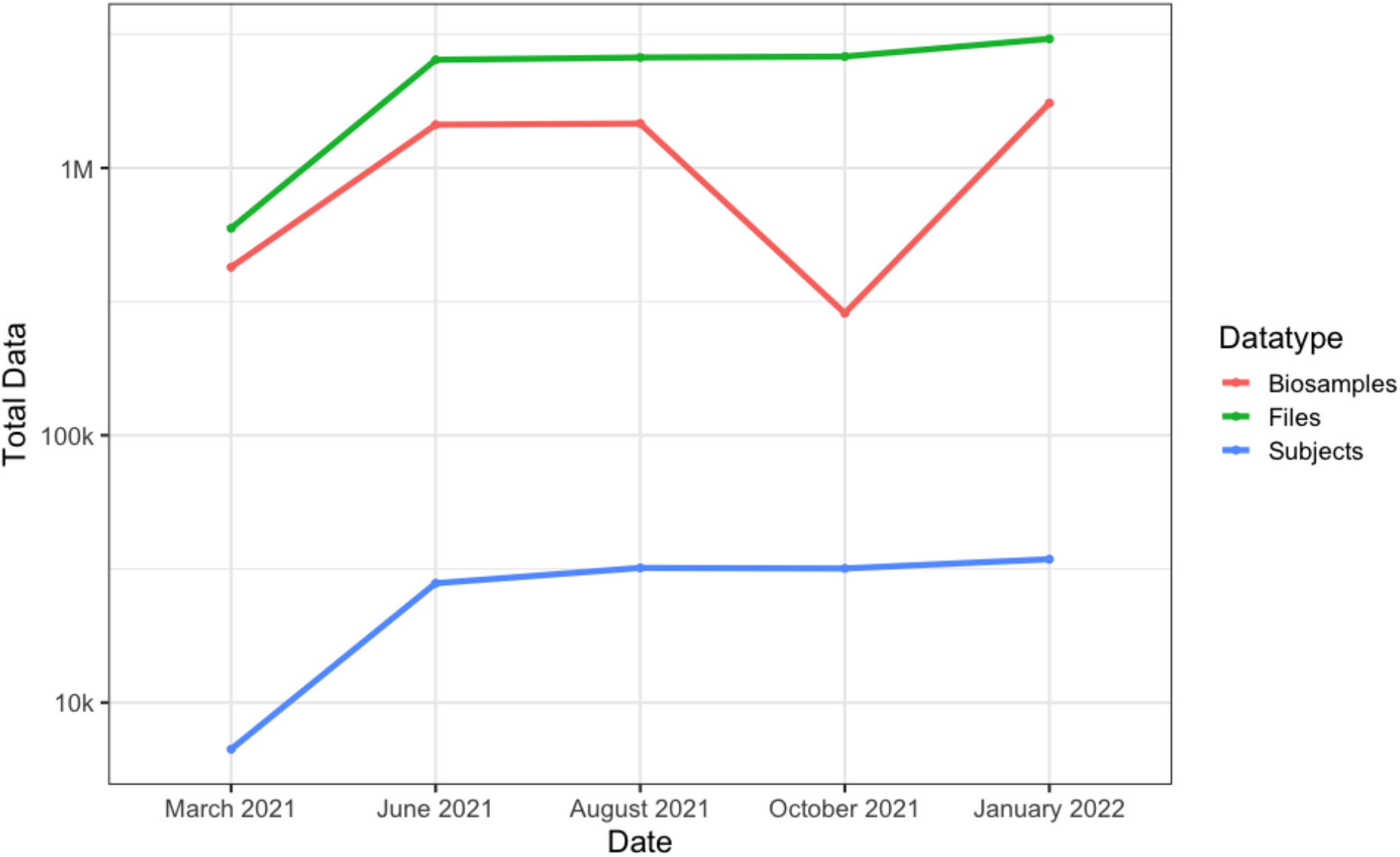
Core data available for search at the CFDE portal over time. The sharp decrease in biosamples in October 2021 is due to replicate cell line data being more appropriately modeled as from a single biosample. Note that the y-axis is exponential, and therefore the increases are quite large, for e.g. the January 2022 release contains nearly half a million (430,405) more files than the October 2021 release.

To enable a DCC to correct errors in a submission and/or to compare different ways of modeling their data, CFDE allows a DCC to submit any number of data packages and to use the CFDE data portal to view each submission in multiple ways. Each public release includes, at most, a single submission from each individual DCC. At each public release date, the most recently approved data package for each DCC is merged into the public catalog and becomes searchable in the portal. If a DCC does not submit and approve a new data package between releases, their current public data package remains in the portal. If a DCC has submitted and approved a new data package, it completely replaces any previous data that was available; submissions cannot currently be incrementally updated.

### The combined C2M2 catalog can be queried via the CFDE portal Web site

Fully processed and approved data packages are merged into the CFDE Portal (https://app.nih-cfde.org/) catalog on a quarterly basis. This catalog is a customized instance of Deriva [43], an asset management platform that provides both a model-driven web interface and a command line interface (deriva-py), with a relational database store that conforms to any model.

The CFDE web interface supports three basic types of search:

1. Search for a specific core entity type (files, biosamples, or subjects) faceted by any number of controlled vocabulary terms
2. Free text search of a single core entity’s controlled vocabulary terms, descriptions, and synonyms
3. Search for core entities associated with a single controlled vocabulary term

The first two search types support use cases where users are interested in finding instances of a specific type of asset (a set of files, a set of biosamples, or a set of subjects). Here, a researcher chooses the type of core entity that they are interested in, and filters from there by using a simple faceted search, faceted search with Boolean operators, or free text search to create a collection of files, biosamples, or subjects. Such searches are especially useful for researchers looking for data similar to that obtained from their own experiments.

Users who are interested in finding data from a given assay, tissue type, disease, or other controlled vocabulary term can use the third search type to filter the CFDE catalog to show all files, biosamples, subjects *and* collections that are associated with that term. This use case allows researchers to quickly assess whether specific data (e.g., mass spectrometry data or psoriasis data) exists in any Common Fund dataset, without needing to specify what type of asset the data is associated with.

Researchers can search the CFDE portal without registration, or can register to access a personalized dashboard page where they can view interactive summary plots, save searches, and build personalized collections of ‘favorite’ items. Registered users can authenticate to the CFDE portal by using a number of identity providers, including eRA Commons.

### Content searchable at the CFDE Portal continues to expand

Following a series of internal prototypes, our first portal release with live data, on March 30, 2021, included submissions from seven Common Fund programs. This first release allowed researchers to search across a combined 594,507 Files, 425,341 Biosamples, and 6,689 Subjects (Figure 4). As of early 2022, the CFDE makes 3,041,978 Files, 1,749,145 Biosamples, and 34,375 Subjects from eleven Common Fund Programs searchable in a single harmonized interface.

The C2M2 is a living standard, and is constantly being expanded to allow new datatypes, new associations, and better ways of describing the underlying data. Over time, DCCs also get better at using the C2M2 to describe their data, a phenomenon clearly visible in the changing number of Biosamples reported over time in Figure 4. In early versions of the portal, DCCs treated cell line replicates as unique Biosamples; as of October, replicate names are collapsed so that the search interface will return all uses of a given named cell line.

### The C2M2 and the CFDE portal are evolving over time to better support search

In the first iteration of the CFDE portal, metadata search was a direct extension of the C2M2 model. However, once DCCs began submitting data packages, it quickly became evident that we needed to provide an extra layer of abstraction between the C2M2 model and users to support intuitive search. The reason was that the terms that DCCs used to describe data often did not correspond to the terms employed by users to search for data.

We use the example of controlled vocabulary anatomy terms to illustrate the difficulty. C2M2 uses UBERON for these terms, and each DCC then maps their local anatomy specifications to UBERON for inclusion in their data package. However, while we expected this mapping process to harmonize anatomy terms across DCCs for easy search, we found in practice that there was no more overlap in term use when the DCCs all used the same vocabulary than when they each used their own idiosyncratic vocabulary. UBERON has, at present, 21,911 unique terms, many describing subtle shades of difference in anatomical structure; in choosing the UBERON term most like their local terminology, DCCs often chose terms at different levels of specificity. We encountered similar results for all controlled vocabularies in our model.

To overcome this barrier to searchability, we have instituted two new practices, one social and one technological. The social solution was to create a working group specifically for dealing with ontology issues, where DCC members can discuss and agree on best practices for choosing ontology terms that are simultaneously a good fit for their data and meaningful to end users. However, even with best practices, there will always be some disagreement on usage. Therefore, we have also created a layer of abstraction in the search portal that allows users to search on higher-level, more general terms under which more specific terms can be grouped through the use of ontology “slims” [44]. For the UBERON anatomy, these “slimmed” search terms are mostly system level names such as nervous system (UBERON:0001016) and connective tissue (UBERON:0002384).

In the CFDE portal, end users can now choose to search by all anatomy terms, only slimmed terms, or both. Similar slim search capabilities are also implemented for the other controlled vocabularies available in C2M2.

### The C2M2 and the CFDE portal are evolving over time to improve user experience

A known challenge for the design of user interfaces to complex systems such as Common Fund data repositories is to ensure that the user intuitively knows how the system should be used. To ensure our portal functionally improves Findability, we have conducted two rounds of usability testing for the CFDE portal interface since launch. These hour-long in-depth interviews were conducted by a professional user experience team [45] to determine both how users currently interact with the portal, as well as how they would like to interact with it. This process revealed a number of assumptions and preferences that we then used to refine the interface to best support user search. In keeping with previous findings, we found that key elements sought on the user interface and portal included: 1) Highly specific terminology or controlled vocabulary; 2) types and extent of data available within the portal; 3) the ability to easily find and complete key tasks; 4) consistent, contextual navigational elements; 5) easily comprehensible data visualizations; 6) the ability to compare tabular data against data visualizations; and 7) dates of data submission. All of these features, as well as many others suggested during these interviews, have been implemented in the existing portal, and will be subject to further refinement from our next testing cohort.

## Discussion

The CFDE portal is a central search solution for locating CF datasets. While a researcher conducting a search sees only a relatively simple user interface, the portal has a complex underlying architecture that relies on the C2M2. The search capabilities of the portal are a visual manifestation of the underlying C2M2, which itself is only useful when populated by data packages. While the CFDE Coordination Center manages and maintains the CFDE infrastructure, we rely on the DCCs to populate the portal with information. This recognition has driven a number of our design decisions around incremental adoption of the C2M2, inclusion of standards, our approach to evolving the C2M2, and our overall approach to federation.

### The C2M2 is designed to support incremental adoption

One specific example of an important evolution that supports improved human engagement on both the data submitter and end user is the addition of slimmed ontology terms; see Results section, above. In brief, after soliciting precise ontology terms from DCCs, we realized that a highly precise terminology hindered discoverability of datasets by end users. We then developed a “slim” ontology that we imposed centrally in order to meet user needs, but did not require DCC engagement. We also created a CFDE-CC working group to allow DCCs to engage in guiding future evolution of the slimmed ontologies. This working group serves as an ongoing interface between the CFDE-CC infrastructure, DCCs, and end users, and supports ongoing engagement around this feature.

C2M2 is intentionally designed to support incremental adoption and use by program participants. In particular, the C2M2 design facilitates the graded introduction of metadata from CF programs into the CFDE system, through submission of data packages with gradually increasing numbers of both entities and associations among entities and controlled vocabulary terms. In turn, this more detailed metadata modeling allows users to conduct more sophisticated searches. Basic C2M2 submissions might consist of basic flat asset inventories, and gradually become well-decorated networks of relationships among resources through a series of small improvements over time.

The C2M2 requires that DCCs meet a fairly sparse set of *minimum* structural benchmarks when building a submission. The general idea is that DCC resource collections can initially be represented quickly (to enable rapid downstream use) via metadata that meets minimal richness requirements -- enough to provide a basic level of harmonization with biomedical experimental metadata coming from other C2M2 sources (DCCs). Over time, DCC data managers can upgrade their C2M2 metadata submissions by adding more detailed descriptive information to their resource records; by elaborating on provenance, timing and other relationships between resources; and by working with CFDE to expand C2M2 itself to better fit models and automation requirements already in production elsewhere.

A minimally compliant submission -- containing just the three required records (Results, above) and no more -- would clearly be of little use. Search capability at the CFDE portal is highly correlated with table richness, and submitters are encouraged (where feasible) to expand and refine the contents of their data package over time. This encouragement is built into the submission system, which displays summary statistics for each draft submission before that submission is integrated with other program submissions for quarterly release.

The ability to submit C2M2 metadata in managed stages of sophistication serves three important purposes. First, it flattens the learning curve for onboarding of DCC data managers by making it possible to create immediately useful submissions with little effort, while encouraging incremental additions over time. Second, it lets DCC data managers test how downstream functionality (e.g., overlapping metadata terms across CFDE) interacts with their C2M2 metadata before investing more heavily in creating more complex C2M2 submissions. Third, it allows *submitters* to provide feedback to CFDE to modify C2M2 in response to submission needs, albeit over longer timelines.

### The C2M2 interfaces seamlessly with existing standards

The world has no shortage of standards, and we have specifically designed our model to leverage mature scientific and technical standards wherever possible. Ultimately a successful metadata model is one that fulfills community needs. In keeping with this philosophy, our initial version of the C2M2 was an evolved version of the DATS model [48], where each data contributor used somewhat different encodings to describe their data, in an ultra flexible system [49]. However, during the in depth interviews conducted during our listening tour, we were able to determine the specific needs of each DCC for modeling their data, as well as learning what metadata was most important to their users. As a result, we completely reimplemented the C2M2 to rely on controlled vocabularies (Table 1) for harmonization, and to require a relatively strict set of metadata tables. This resulted in a somewhat less flexible model, but one that is still more than nimble enough to meet the needs of the CFDE community, while also supporting the types of faceted search that biomedical researchers prefer.

The C2M2 mission is to operationalize and anchor our guiding principles of data stewardship -- findability, accessibility, interoperability, and reusability (FAIR) -- all of which are enhanced by the integration of established standards directly into the model framework. Findability is streamlined by the use of common terms to describe scientific concepts: aggregating data according to harmonized and universal descriptive metadata helps users find information and enhances discoverability of relevant related data. Accessibility benefits from integration of technical standards allowing users uniform access to heterogeneous data sources without needing to use multiple bespoke access interfaces. Interoperability is defined by how well data flows interact with other information systems: adoption of technical interface standards directly determines how interoperable any system will be. Reusability depends both on persistence of data over time (encouraged directly in C2M2 by rules governing persistent identifiers) and on the implementation of conceptual standards defining meaning and context (so future users can properly explore the data for their own purposes).

By design, the C2M2 will be amended and extended over time to include additional metadata and relationships, including new community standards, so it can flexibly grow to support any biomedical metadata type. Future work to improve global discoverability of CFDE Portal resources may for example include integrations with harmonizing efforts like schema.org [50] or bioschemas [51]. Technologies like these aim both to increase data accessibility via common query interfaces and to improve search engine visibility for indexed resources and datasets.

All of the C2M2 modeling decisions were made to strike a balance between ease of harmonization, and ease of user search. The C2M2 supports two distinct identifiers for each entity, a (required) C2M2-specific ID and also an (optional) persistent ID which is globally resolvable. Required C2M2 IDs can be automatically generated from local DCC identifiers, avoiding the need to mint new IDs before submitting to the CFDE portal and thereby reducing cost and complexity for DCCs preparing C2M2 metadata submissions. Optional persistent IDs can be used to give users direct access (via extramural protocols and APIs) to further metadata describing experimental resources of interest, including direct or programmatic download access to data files indexed by CFDE. Persistent IDs constitute a critical element of the C2M2 framework: they facilitate structured, stable, and reliable access to research information housed outside the CFDE portal.

### Use of the C2M2 is supported by extensive documentation

Participating DCCs create their submissions by mapping their internal data model to the C2M2; depending on the complexity of their data, this mapping can be a difficult task. To support DCCs in creating their data packages, we provide full technical documentation [52], a more novice friendly wiki [53], a bug and request tracker [54], and a full service helpdesk. Our submission system and portal review system also provide detailed error messages for invalid data packages.

### The C2M2 is constantly evolving and expanding

The purpose of the C2M2 is to facilitate metadata harmonization: thus, wherever possible, it should represent legitimately comparable entities in standard ways without compromising meaning, context, or accuracy. Where it may be useful to weaken precision to preserve search recall, slims ensure that the underlying metadata remain accessible. Importantly, the evolution of the C2M2 is an ongoing process.

Most DCCs already use some internal metadata model for their own curation operations. C2M2 representation of similar but distinct packages of important information, taken from multiple independently developed custom DCC metadata systems (e.g., metadata about people and organizations, data provenance, experimental protocols, or detailed event sequences) requires ongoing, iterative, case-based design and consensus-driven decision-making, coordinated across multiple research groups. Design and decision-making in such contexts requires long-term planning, testing, and execution. CFDE is committed to handling new metadata that are difficult to integrate and harmonize by the creation of generalizable, well-defined extensions to C2M2 if possible, and by pruning (at least in the short term) if not.

We aim with the flexible C2M2 design to split the difference between the ease of evolution inherent in a simple model and the operational power provided to downstream applications by more detailed but difficult-to-maintain extended frameworks. This flexibility is also intended to address the needs of different DCCs that inevitably operate at widely different scales of data complexity or funding level as well as organization life-cycle phases, research scope, etc. DCCs with advanced, operationalized metadata modeling systems of their own should not encounter arbitrary barriers to C2M2 support for more extensive relational modeling of their metadata if they want it; newer or smaller DCCs, by contrast, may not have enough readily-available information to feasibly describe their experimental resources beyond giving basic asset lists and project attributions. By committing to developing modular C2M2 extensions for the most advanced DCC metadata, while also offering simpler but well-structured model options for simpler data (that are, furthermore, already harmonized across C2M2 metadata from other DCCs) we aim to minimize barriers to rapid entry into the C2M2 ecosystem and its downstream applications. This approach both allows us to meet the needs of an ever expanding group of stakeholders, and makes C2M2 an ideal framework for other consortia to adopt for their own data curation needs.

## Supporting information

SuppFig1

SuppText

## Funding

NIH Common Fund OT3OD025459-01 for the CFDE Coordinating Center

## Data Availability

Full C2M2 Documentation: https://docs.nih-cfde.org/en/latest/c2m2/draft-C2M2_specification/ C2M2 Supplemental Wiki: https://github.com/nih-cfde/published-documentation/wiki

C2M2 Submission System Documentation: https://docs.nih-cfde.org/en/latest/cfde-submit/docs/

**Portal code:**

CFDE specific Deriva fork: https://github.com/nih-cfde/cfde-deriva

Portal UI customization: https://github.com/nih-cfde/dashboard

**Submission code:**

CFDE specific Globus flow fork: https://github.com/nih-cfde/cfde-submit

CFDE specific Globus action provider fork: https://github.com/nih-cfde/deriva-action-provider

## Notes

https://app.nih-cfde.org/

### Competing Interest Statement

The authors have declared no competing interest.

### Summary of Updates

Replace Supplemental Figure 1. Previous had bad figure cropping.

