## Supplementary material for "Making Common Fund data more findable: Catalyzing a Data Ecosystem": SuppFig1

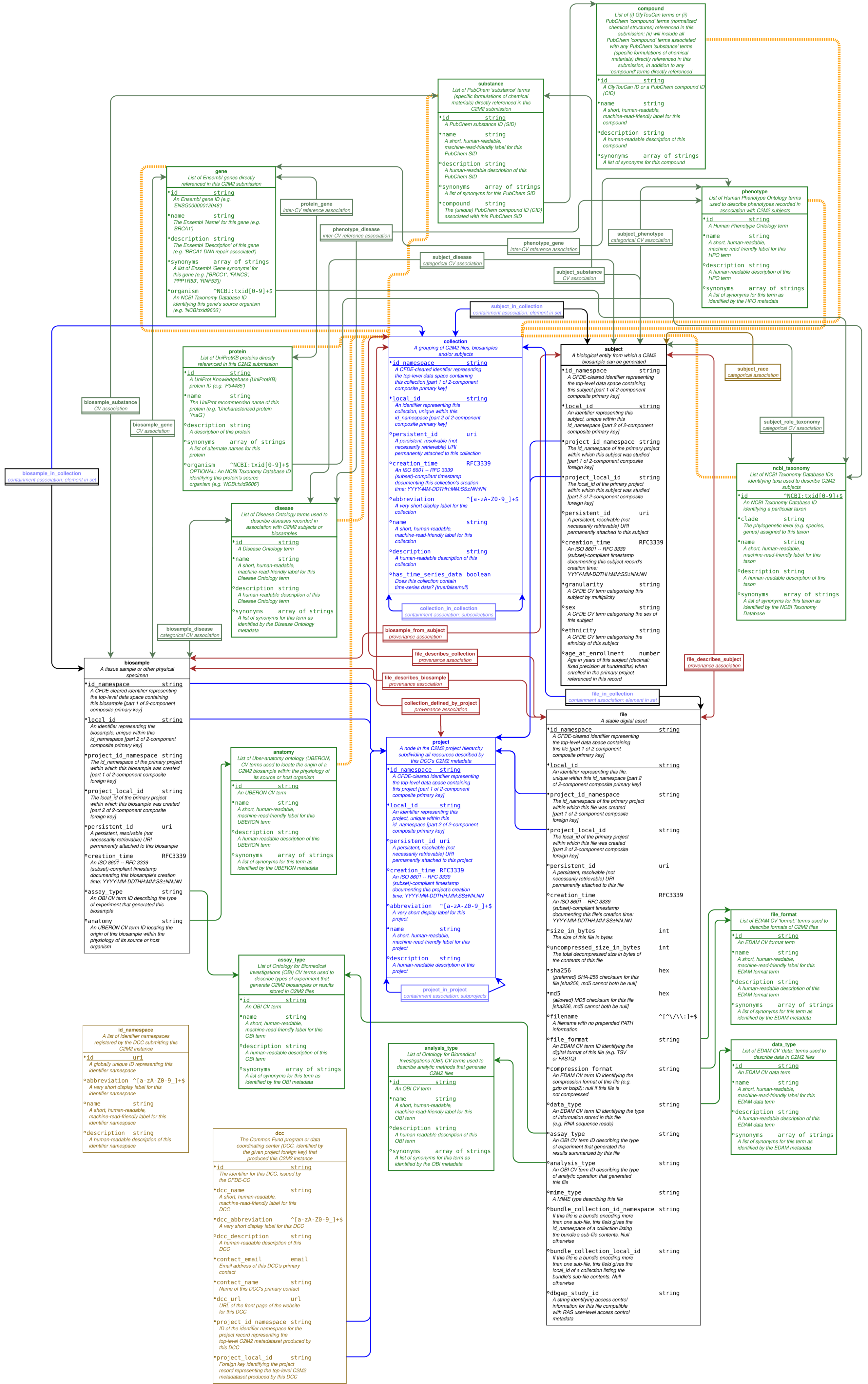

compound
List of (i) GlyTouCan terms or (ii) PubChem 'compound' terms (normalized chemical structures) referenced in this submission; (ii) will include all PubChem 'compound' terms associated with any PubChem 'substance' terms (specific formulations of chemical materials) directly referenced in this submission, in addition to any 'compound' terms directly referenced
id string
A GlyTouCan ID or a PubChem compound ID (CID)
name string
A short, human-readable, machine-read-friendly label for this compound
description string
A human-readable description of this compound
synonyms array of strings
A list of synonyms for this compound
substance
List of PubChem 'substance' terms (specific formulations of chemical materials) directly referenced in this C2M2 submission
id string
A PubChem substance ID (SID)
name string
A short, human-readable, machine-read-friendly label for this PubChem SID
description string
A human-readable description of this PubChem SID
synonyms array of strings
A list of synonyms for this PubChem SID
compound string
The (unique) PubChem compound ID (CID) associated with this PubChem SID
gene
List of Ensembl genes directly referenced in this C2M2 submission
id string
An Ensembl gene ID (e.g. 'ENSG00000012048')
name string
The Ensembl 'Name' for this gene (e.g. 'BRCA1')
description string
The Ensembl 'Description' of this gene (e.g. 'BRCA1 DNA repair associated')
synonyms array of strings
A list of Ensembl 'Gene synonyms' for this gene (e.g. ['BRCC1', 'FANCS', 'PPP1R53', 'RNF53'])
organism ^NCBI:txid[0-9]+$
An NCBI Taxonomy Database ID identifying this gene's source organism (e.g. 'NCBI:txid9606')
protein_gene
inter-CV reference association
phenotype_disease
inter-CV reference association
phenotype_gene
inter-CV reference association
subject_disease
categorical CV association
subject_phenotype
categorical CV association
subject_substance
CV association
phenotype
List of Human Phenotype Ontology terms used to describe phenotypes recorded in association with C2M2 subjects
id string
A Human Phenotype Ontology term
name string
A short, human-readable, machine-read-friendly label for this HPO term
description string
A human-readable description of this HPO term
synonyms array of strings
A list of synonyms for this term as identified by the HPO metadata
subject_in_collection
containment association: element in set
protein
List of UniProtKB proteins directly referenced in this C2M2 submission
id string
A UniProt Knowledgebase (UniProtKB) protein ID (e.g. 'P94485')
name string
The UniProt recommended name of this protein (e.g. 'Uncharacterized protein YnaG')
description string
A description of this protein
synonyms array of strings
A list of alternate names for this protein
organism ^NCBI:txid[0-9]+$
OPTIONAL: An NCBI Taxonomy Database ID identifying this protein's source organism (e.g. 'NCBI:txid9606')
collection
A grouping of C2M2 files, biosamples and/or subjects
id_namespace string
A CFDE-cleared identifier representing the top-level data space containing this collection [part 1 of 2-component composite primary key]
local_id string
An identifier representing this collection, unique within this id_namespace [part 2 of 2-component composite primary key]
persistent_id uri
A persistent, resolvable (not necessarily retrievable) URI permanently attached to this collection
creation_time RFC3339
An ISO 8601 -- RFC 3339 (subset)-compliant timestamp documenting this collection's creation time: YYYY-MM-DDTHH:MM:SS±NN:NN
abbreviation ^[a-zA-Z0-9_]+$
A very short display label for this collection
name string
A short, human-readable, machine-read-friendly label for this collection
description string
A human-readable description of this collection
has_time_series_data boolean
Does this collection contain time-series data? (true/false/null)
collection_in_collection
containment association: subcollections
subject
A biological entity from which a C2M2 biosample can be generated
id_namespace string
A CFDE-cleared identifier representing the top-level data space containing this subject [part 1 of 2-component composite primary key]
local_id string
An identifier representing this subject, unique within this id_namespace [part 2 of 2-component composite primary key]
project_id_namespace string
The id_namespace of the primary project within which this subject was studied [part 1 of 2-component composite foreign key]
project_local_id string
The local_id of the primary project within which this subject was studied [part 2 of 2-component composite foreign key]
persistent_id uri
A persistent, resolvable (not necessarily retrievable) URI permanently attached to this subject
creation_time RFC3339
An ISO 8601 -- RFC 3339 (subset)-compliant timestamp documenting this subject record's creation time: YYYY-MM-DDTHH:MM:SS±NN:NN
granularity string
A CFDE CV term categorizing this subject by multiplicity
sex string
A CFDE CV term categorizing the sex of this subject
ethnicity string
A CFDE CV term categorizing the ethnicity of this subject
age_at_enrollment number
Age in years of this subject (decimal: fixed precision at hundredths) when enrolled in the primary project referenced in this record
subject_race
categorical association
subject_role_taxonomy
categorical CV association
ncbi_taxonomy
List of NCBI Taxonomy Database IDs identifying taxa used to describe C2M2 subjects
id ^NCBI:txid[0-9]+$
An NCBI Taxonomy Database ID identifying a particular taxon
clade string
The phylogenetic level (e.g. species, genus) assigned to this taxon
name string
A short, human-readable, machine-read-friendly label for this taxon
description string
A human-readable description of this taxon
synonyms array of strings
A list of synonyms for this taxon as identified by the NCBI Taxonomy Database
biosample_substance
CV association
biosample_gene
CV association
biosample_in_collection
containment association: element in set
disease
List of Disease Ontology terms used to describe diseases recorded in association with C2M2 subjects or biosamples
id string
A Disease Ontology term
name string
A short, human-readable, machine-read-friendly label for this Disease Ontology term
description string
A human-readable description of this Disease Ontology term
synonyms array of strings
A list of synonyms for this term as identified by the Disease Ontology metadata
biosample_disease
categorical CV association
biosample_from_subject
provenance association
file_describes_collection
provenance association
file_describes_subject
provenance association
file_describes_biosample
provenance association
collection_defined_by_project
provenance association
file_in_collection
containment association: element in set
biosample
A tissue sample or other physical specimen
id_namespace string
A CFDE-cleared identifier representing the top-level data space containing this biosample [part 1 of 2-component composite primary key]
local_id string
An identifier representing this biosample, unique within this id_namespace [part 2 of 2-component composite primary key]
project_id_namespace string
The id_namespace of the primary project within which this biosample was created [part 1 of 2-component composite foreign key]
project_local_id string
The local_id of the primary project within which this biosample was created [part 2 of 2-component composite foreign key]
persistent_id uri
A persistent, resolvable (not necessarily retrievable) URI permanently attached to this biosample
creation_time RFC3339
An ISO 8601 -- RFC 3339 (subset)-compliant timestamp documenting this biosample's creation time: YYYY-MM-DDTHH:MM:SS±NN:NN
assay_type string
An OBI CV term ID describing the type of experiment that generated this biosample
anatomy string
An UBERON CV term ID locating the origin of this biosample within the physiology of its source or host organism
anatomy
List of Uber-anatomy ontology (UBERON) CV terms used to locate the origin of a C2M2 biosample within the physiology of its source or host organism
id string
An UBERON CV term
name string
A short, human-readable, machine-read-friendly label for this UBERON term
description string
A human-readable description of this UBERON term
synonyms array of strings
A list of synonyms for this term as identified by the UBERON metadata
project
A node in the C2M2 project hierarchy subdividing all resources described by this DCC's C2M2 metadata
id_namespace string
A CFDE-cleared identifier representing the top-level data space containing this project [part 1 of 2-component composite primary key]
local_id string
An identifier representing this project, unique within this id_namespace [part 2 of 2-component composite primary key]
persistent_id uri
A persistent, resolvable (not necessarily retrievable) URI permanently attached to this project
creation_time RFC3339
An ISO 8601 -- RFC 3339 (subset)-compliant timestamp documenting this project's creation time: YYYY-MM-DDTHH:MM:SS±NN:NN
abbreviation ^[a-zA-Z0-9_]+$
A very short display label for this project
name string
A short, human-readable, machine-read-friendly label for this project
description string
A human-readable description of this project
project_in_project
containment association: subprojects
file
A stable digital asset
id_namespace string
A CFDE-cleared identifier representing the top-level data space containing this file [part 1 of 2-component composite primary key]
local_id string
An identifier representing this file, unique within this id_namespace [part 2 of 2-component composite primary key]
project_id_namespace string
The id_namespace of the primary project within which this file was created [part 1 of 2-component composite foreign key]
project_local_id string
The local_id of the primary project within which this file was created [part 2 of 2-component composite foreign key]
persistent_id uri
A persistent, resolvable (not necessarily retrievable) URI permanently attached to this file
creation_time RFC3339
An ISO 8601 -- RFC 3339 (subset)-compliant timestamp documenting this file's creation time: YYYY-MM-DDTHH:MM:SS±NN:NN
size_in_bytes int
The size of this file in bytes
uncompressed_size_in_bytes int
The total decompressed size in bytes of the contents of this file
sha256 hex
(preferred) SHA-256 checksum for this file [sha256, md5 cannot both be null]
md5 hex
(allowed) MD5 checksum for this file [sha256, md5 cannot both be null]
filename ^[^\/\\:]+$
A filename with no prepended PATH information
file_format string
An EDAM CV term ID identifying the digital format of this file (e.g. TSV or FASTQ)
compression_format string
An EDAM CV term ID identifying the compression format of this file (e.g. gzip or bzip2): null if this file is not compressed
data_type string
An EDAM CV term ID identifying the type of information stored in this file (e.g. RNA sequence reads)
assay_type string
An OBI CV term ID describing the type of experiment that generated the results summarized by this file
analysis_type string
An OBI CV term ID describing the type of analytic operation that generated this file
mime_type string
A MIME type describing this file
bundle_collection_id_namespace string
If this file is a bundle encoding more than one sub-file, this field gives the id_namespace of a collection listing the bundle's sub-file contents. Null otherwise
bundle_collection_local_id string
If this file is a bundle encoding more than one sub-file, this field gives the local_id of a collection listing the bundle's sub-file contents. Null otherwise
dbgap_study_id string
A string identifying access control information for this file compatible with RAS user-level access control metadata
file_format
List of EDAM CV 'format:' terms used to describe formats of C2M2 files
id string
An EDAM CV format term
name string
A short, human-readable, machine-read-friendly label for this EDAM format term
description string
A human-readable description of this EDAM format term
synonyms array of strings
A list of synonyms for this term as identified by the EDAM metadata
data_type
List of EDAM CV 'data:' terms used to describe data in C2M2 files
id string
An EDAM CV data term
name string
A short, human-readable, machine-read-friendly label for this EDAM data term
description string
A human-readable description of this EDAM data term
synonyms array of strings
A list of synonyms for this term as identified by the EDAM metadata
assay_type
List of Ontology for Biomedical Investigations (OBI) CV terms used to describe types of experiment that generate C2M2 biosamples or results stored in C2M2 files
id string
An OBI CV term
name string
A short, human-readable, machine-read-friendly label for this OBI term
description string
A human-readable description of this OBI term
synonyms array of strings
A list of synonyms for this term as identified by the OBI metadata
analysis_type
List of Ontology for Biomedical Investigations (OBI) CV terms used to describe analytic methods that generate C2M2 files
id string
An OBI CV term
name string
A short, human-readable, machine-read-friendly label for this OBI term
description string
A human-readable description of this OBI term
synonyms array of strings
A list of synonyms for this term as identified by the OBI metadata
id_namespace
A list of identifier namespaces registered by the DCC submitting this C2M2 instance
id uri
A globally unique ID representing this identifier namespace
abbreviation ^[a-zA-Z0-9_]+$
A very short display label for this identifier namespace
name string
A short, human-readable, machine-read-friendly label for this identifier namespace
description string
A human-readable description of this identifier namespace
dcc
The Common Fund program or data coordinating center (DCC, identified by the given project foreign key) that produced this C2M2 instance
id string
The identifier for this DCC, issued by the CFDE-CC
dcc_name string
A short, human-readable, machine-read-friendly label for this DCC
dcc_abbreviation ^[a-zA-Z0-9_]+$
A very short display label for this DCC
dcc_description string
A human-readable description of this DCC
contact_email email
Email address of this DCC's primary contact
contact_name string
Name of this DCC's primary contact
dcc_url url
URL of the front page of the website for this DCC
project_id_namespace string
ID of the identifier namespace for the project record representing the top-level C2M2 metadataset produced by this DCC
project_local_id string
Foreign key identifying the project record representing the top-level C2M2 metadataset produced by this DCC

Figure Supplement-1. An **entity-relationship (ER) diagram** describing C2M2. Entities (things) are drawn as full tables: boxes with descriptions of named fields. **Associations** (relationships between entities) are named inside small boxes: arrows are drawn connecting each association with the entities that participate in the relationship that the association represents. Color key:

■ Black: **Core entities** (basic experimental resources): `file`, `biosample` and `subject`

■ Dark red: **Association relationships** between entities

■ Blue: **Container entities** (`project` and `collection`) and their containment relationships

■ Green: **Term entities and associations** for all standardized controlled-vocabulary terms submitted as C2M2 annotation metadata, plus extra descriptive information to facilitate user searching and web displays

■ Gold: **Administrative entities** giving technical contact information for submitters and describing DCC-controlled identifier namespaces

■ Yellow: association tables **linking `collections` to CVs** describing anatomy, compounds, substances, proteins, genes, diseases and phenotypes
