## Supplementary material for "Making Common Fund data more findable: Catalyzing a Data Ecosystem": SuppText

### C2M2 Persistent IDs

C2M2 metadata are managed and curated by Common Fund DCCs to standardize and stabilize it for future research use. CFDE's explicit mission for C2M2 is to create an information archive that can usefully serve researchers who work without guaranteed access to follow-up information, including (among other scenarios) for future work done after the funding lifecycle of each managing DCC has ended.

To be used as a C2M2 `persistent_id`, an ID

1. represents an explicit commitment by the managing DCC that the attachment of the ID to the resource it represents is **permanent and final**
2. must be a format-compliant URI or a compact identifier, where the protocol (the "scheme" or "prefix") specified in the ID is registered with at least one of the following (see the given lists for examples of URIs and compact identifiers)
    - IANA (list of registered schemes)
        - scheme used must be assigned either "Permanent" or "Provisional" status
    - Identifiers.org (list of registered prefixes)
    - N2T (Name-To-Thing) (list of registered prefixes)
3. if representing a `file` and used as a `persistent_id`, **cannot** be a direct-download URL for that `file`: it must instead be an identifier permanently attached to the `file` and only **indirectly resolvable** (through the scheme or prefix specified within the ID) to the `file` itself

These requirements constitute a minimal set of rules to ensure that C2M2 resources can be stably cited in scientific literature and automatically reused in future research. Clearly, though, the production and maintenance of `persistent_ids` represents a substantial investment of time, thought, and effort, and we also emphasize that not every C2M2 resource record that *can* receive a `persistent_id` will necessarily ever *need* one. These IDs -- while representing a gold standard for stability and long-term access -- are **strictly optional**. DCCs should also note that without `persistent_ids`, digital file assets represented in C2M2 will serve *only* as inventory items and annotated search results: permanent, indirected `persistent_ids` are required in order to enable *any* automated interoperability between actual data files referenced by C2M2 records and external software systems (including direct download access to files).

Since `persistent_id` is always optional, C2M2 provides a separate structure to provide for universal identification: the basic **C2M2 ID** is a two-part label comprised of a prefix (`id_namespace`) and a suffix (`local_id`) which, concatenated, make up the ID. C2M2 IDs fall into categories described by three main cases:

**[1]** A `persistent_id` already exists for the object being named.

- if the `persistent_id` is a URI, then that URI should be split to form a C2M2 ID (see the URI reference for precise definitions of terms like "scheme" and "path" in this context):
    - `id_namespace` (prefix): `scheme://authority/`
    - `local_id` (suffix): `path`
    - Example: an SRA accession URI `https://www.ncbi.nlm.nih.gov/sra/SRX000007` stored in C2M2 as a `persistent_id` would be split, to form a corresponding C2M2 ID, into
        - an `id_namespace` prefix of `https://www.ncbi.nlm.nih.gov/sra/`
        - and a `local_id` suffix of `SRX000007`
- if the existing `persistent_id` is not a URI but instead is a compact identifier, it should be split similarly, with the details determined according to the particular format specification for the prefix being used: a scheme label and a reference to the issuing or owning authority (plus a delimiter) should constitute the `id_namespace` prefix, and the ID of the particular thing being referenced should be stored in the `local_id` suffix.
    - Example: the DOI compact identifier `doi:10.1006/jmbi.1998.2354` would be split into
        - an `id_namespace` prefix of `doi:10.1006/`, specifying the identifier type (`doi`) and the registered owner of the object (`10.1006`)
        - and a `local_id` suffix of `jmbi.1998.2354`

**[2]** A DCC already uses URIs to identify things that correspond to C2M2 entities (`files`, `biosamples`, etc.), but those URIs don't meet all the criteria to be C2M2 `persistent_ids` (e.g. they're not guaranteed to be permanent). Such URIs can still be split into an `id_namespace` prefix (containing a reference to the controlling authority, e.g. the DCC or one of its organizational data sources) and a `local_id` suffix (describing the object being identified) to form a C2M2 ID. (For records with IDs built like this, `persistent_id` would be left blank.)

**[3]** A DCC only has local identifiers for such entities. In this case, each local identifier will be the corresponding C2M2 `local_id` suffix (sanitized as necessary for URI safety), and the `id_namespace` prefix can be constructed according to the 'tag' URI proposal.

- Example: The tag-URI-based `id_namespace/local_id` C2M2 ID for a C2M2 `biosample` record representing Sample A-867-5309 at the Flerbiger's Disease Project (FDP) -- a non-permanent, strictly local sample ID assigned by the FDP for their C2M2 submission built at the end of the first quarter of 2021 -- might be (an email address would also work in place of 'flerbiger.org' below)
    - `id_namespace`: `tag:flerbiger.org,2021-03-31:`
    - `local_id`: `A-867-5309`
